# Spatial memory formation requires netrin-1 expression by neurons in the adult mammalian brain

**DOI:** 10.1101/490680

**Authors:** Edwin W. Wong, Stephen D. Glasgow, Lianne J. Trigiani, Daryan Chitsaz, Vladmir Rymar, Abbas Sadikot, Edward S. Ruthazer, Edith Hamel, Timothy E. Kennedy

**Author notes:** indicates equal contribution (Edwin W. Wong, Stephen D. Glasgow). Corresponding Author and Lead Contact: Timothy E. Kennedy, Ph.D., co-Director, McGill Program in Neuroengineering, Professor, Department of Neurology and Neurosurgery, Montreal Neurological Institute, McGill University, 3801 University Avenue, Montreal, Quebec, Canada, H3A 2B4.

## Abstract

Netrin-1 was initially characterized as an axon guidance molecule that is essential for normal embryonic neural development; however, many types of neurons continue to express netrin-1 in the post-natal and adult mammalian brain. Netrin-1 and the netrin receptor DCC are both enriched at synapses. In the adult hippocampus, activity-dependent secretion of netrin-1 by neurons potentiates glutamatergic synapse function, and is critical for long-term potentiation, an experimental cellular model of learning and memory. Here, we assessed the impact of neuronal expression of netrin-1 in the adult brain on behavior using tests of learning and memory. We show that adult mice exhibit impaired spatial memory following conditional deletion of netrin-1 from glutamatergic neurons in the hippocampus and neocortex. Further, we provide evidence that mice with conditional deletion of netrin-1 do not display aberrant anxiety-like phenotypes and show a reduction in self-grooming behaviour. These findings reveal a critical role for netrin-1 expressed by neurons in the regulation of spatial memory formation.

## Introduction

Secreted chemotropic guidance cues direct axon extension during embryogenesis in the developing nervous system, yet after axon guidance is complete, many of these cues continue to be expressed by neurons and glia in the adult. Expression of guidance cues and their receptors by neurons suggests that these proteins may contribute to mature neuronal function, including synaptic plasticity underlying learning and memory (Shen and Cowan 2010). Memory consolidation is thought to involve the modification of synaptic structure and function (Roberts et al. 2010), though how guidance cues may contribute to these changes remains unclear.

Netrin-1, a canonical secreted guidance cue, is a laminin-related protein that directs axon extension and promotes synapse formation during early development (Kennedy et al. 1994; Serafini et al. 1994; Goldman et al. 2013). The netrin receptor, deleted in colorectal cancer (DCC) (Keino-Masu et al. 1996) triggers increases in intracellular calcium, activation of RhoGTPases such as Cdc42 and Rac1, and regulates local protein synthesis (Kim and Martin, 2015; Lai Wing Sun et al., 2011). Netrin-1 and DCC are highly enriched at synapses in the mature mammalian brain and DCC co-fractionates with detergent-resistant components of the post-synaptic density (Horn et al. 2013). We have recently reported that netrin-1 is released at synaptic sites in response to N-methyl-D-aspartate glutamate receptor (NMDAR) activation and is critical for expression of long-term potentiation at hippocampal Schaffer-collateral synapses, an experimental model of synaptic plasticity in the adult brain (Glasgow et al. 2018). Further, application of exogenous netrin-1 is sufficient to trigger insertion of GluA1-containing α-amino-3-hydroxy-5-methyl-4-isoxazolepropionic acid glutamate receptors (AMPARs). Together, these findings indicate that netrin-1 participates in activity-dependent plasticity at Schaffer-collateral synapses, that netrin-1 is secreted by neurons in response to activity, and that netrin-1 is sufficient to evoke lasting synaptic potentiation (Glasgow et al. 2018). Here we report that conditional deletion of netrin-1 from principal excitatory neurons results in deficits in hippocampal-dependent spatial memory, demonstrating that netrin-1 critically regulates memory processes underlying spatial cognition.

## Results

### Selective deletion of netrin-1 from forebrain glutamatergic neurons impairs spatial memory

We have recently reported that neuronal expression of netrin-1 in principal excitatory neurons impairs long-term potentiation in the adult hippocampus, suggesting that netrin-1 may be necessary for spatial memory formation (Glasgow et al. 2018). To test the hypothesis that netrin-1 made by glutamatergic neurons in the forebrain contributes to memory formation, we tested CaMKII-Cre/NTN1^*f/f*^ (NTN1 cKO) and wild-type age-matched littermate controls in the hippocampus-dependent Morris water maze (MWM) task (Morris et al. 1982). For the first three days, the “visible” phase, mice were trained to swim to a visible platform cued by a marked object in the maze and spatial cues within the room. The “hidden” phase followed from days 4 to 8, during which spatial cues in the room were switched and the mice were challenged to locate a hidden platform placed in a different quadrant of the maze with the visible cue removed. All animals performed similarly with regard to escape latency to reach the platform, indicating intact sensory and motor functions (Figure 1A). The observed improvement in performance across training days is consistent with the formation of a cognitive spatial map (Brody and Holtzman 2006). Twenty-four hours following the final training session (Day 8), the platform was removed. We observed no significant differences in the swimming speed of NTN1 cKO and *cre-*negative wild-type littermate mice (Figure 1B). However, adult NTN1 cKO mice made significantly fewer crosses over the location of the removed platform during the probe trial, spent less time within the target quadrant, and travelled less distance within the target quadrant compared to control wild-type littermate mice (Figures 1C-E). These findings provide evidence that neuronal expression of netrin-1 is critical for spatial memory consolidation and precision.

**Figure 1.**
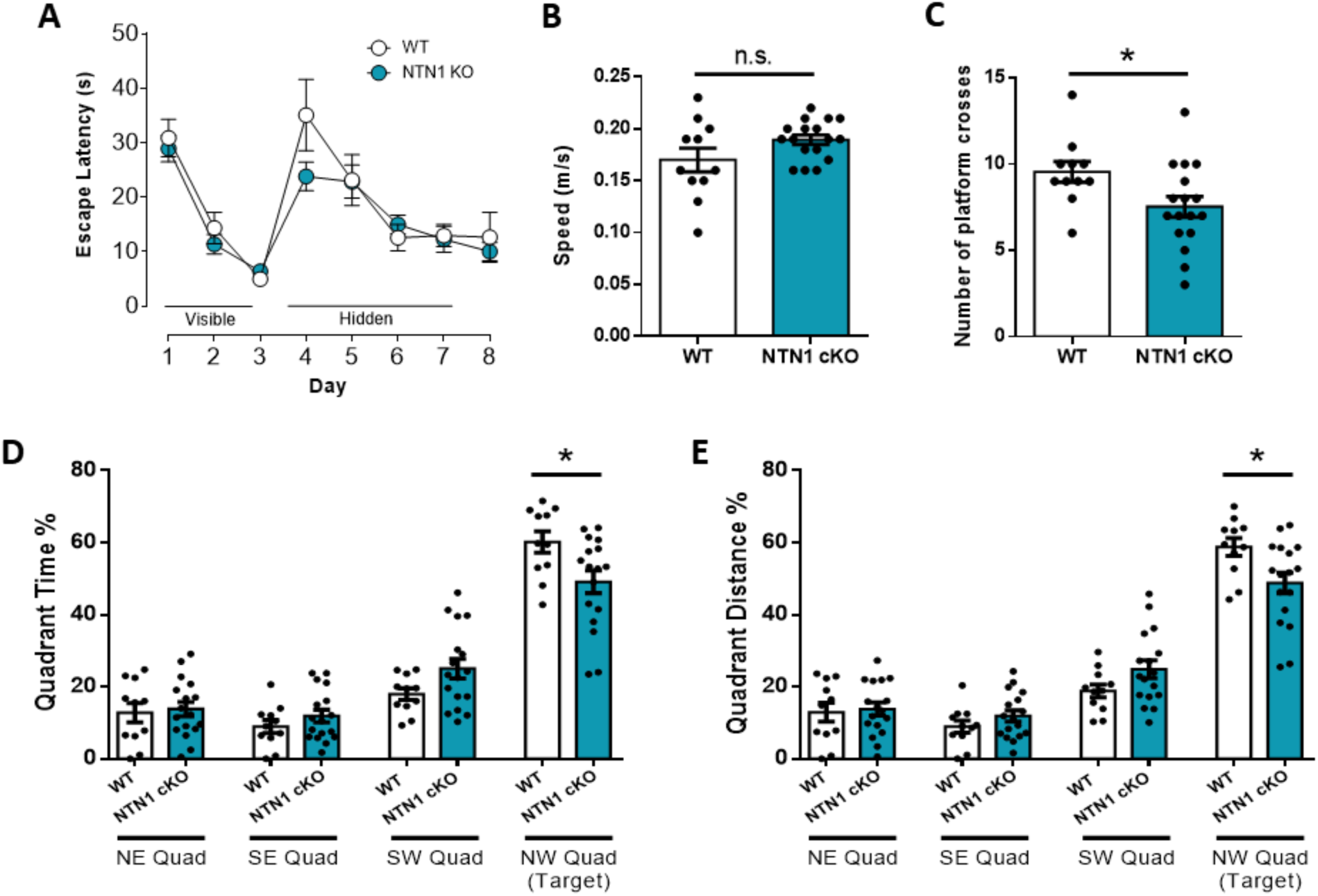
Mice with conditional deletion of netrin-1 from glutamatergic neurons in the forebrain exhibit spatial memory deficits in the Morris water maze. (**A**) Both wild-type and NTN1 cKO mice showed no differences in performance during the training phases (WT: *n* = 11, NTN1 cKO: *n* = 17; two-way repeated measure ANOVA, Bonferroni’s post-hoc). (B-E) No differences were observed in speed between genotypes (**B**; WT: 0.1700 ± 0.011, NTN1 cKO: 0.1894 ± 0.0046, *p* = 0.083). NTN1 cKO mice (blue bars) made fewer passes over platform location location (**C**; WT: 9.545 ± 0.593, NTN1 cKO: 7.529 ± 0.59, *p* = 0.031), spent proportionally less time (**D**; WT: 58.75 ± 2.47, NTN1 cKO: 48.82 ± 2.89, *p* = 0.022), and traveled significantly less distance (**E**; WT: 60.18 ± 2.93, NTN1 cKO: 49.11 ± 3.13, *p* = 0.023) in the target quadrant compared to control mice (white bars). All comparisons were performed with two-tailed independent t-test where *\*p* < 0.05. Data are shown as mean ± SEM.

While the impairments described above suggest a deficit in spatial memory in NTN1 cKO mice, performance in the MWM can also engage the encoding and retrieval of emotionally-aversive training events (D’Hooge and De Deyn 2001). To test whether neuronal netrin-1 expression is necessary for hippocampal-dependent spatial memory consolidation, we tested NTN1 cKO and age-matched control littermate mice using the novel object place recognition (NOPR) task (Figure 2A). We observed no significant differences in total exploration time between NTN1 cKO and age-matched wild-type littermate control mice (Figure 2B). In contrast, we observed significant decreases in discrimination index and reduced investigative ratios in NTN1 cKO mice compared to control littermates (Figures 2C-2D). Control littermates also showed an expected higher interaction count for the novel placed object compared to the unmoved, familiar placed object during the Choice Phase, while NTN1 cKO mice showed no differences between the two objects (Figure 2E). Together, these findings strongly implicate a critical role for neuronal expression of netrin-1 in spatial memory.

**Figure 2.**
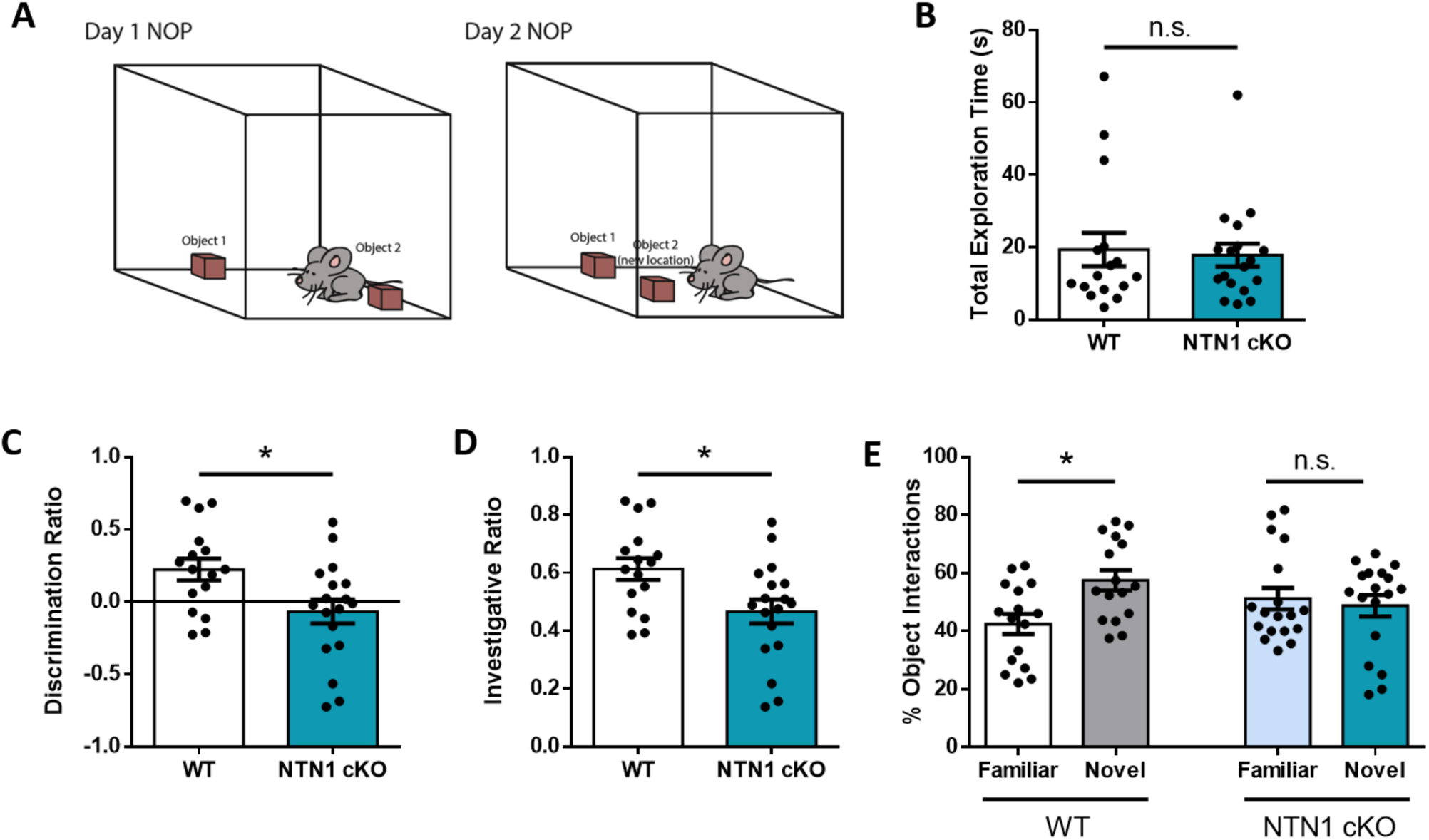
NTN1 cKO mice exhibit spatial memory deficits in the novel object place recognition (NOPR) test. (**A**) Schematic of the NOPR test. (**B**) No significant differences were observed in the total exploration time for either object (WT: *n* = 16, 19.36 ± 4.58, NTN1 cKO: *n* = 18, 17.86 ± 3.17, *p* = 0.79). (**C**-**D**) NTN1 cKO scored significant less on the discrimination ratio (**C**; WT: 0.223 ± 0.074, NTN1 cKO: −0.067 ± 0.083, *p* = 0.014) and investigative ratio (**D**;WT: 0.6137 ± 0.037, NTN1 cKO: 0.4664 ± 0.041, *p* = 0.013) during the Choice Phase. (**E**) Wild-type mice displayed significantly more interactions with the novel placed object compared to the familiar object but not NTN1 cKO animals (WT Familiar: 42.46 ± 3.47, WT Novel: 57.54 ± 3.47, NTN1 cKO Familiar: 51.21 ± 3.71, NTN1 cKO Novel: 48.79 ± 3.71, Genotype × Choice interaction: *F*_3, 64_ = 2.82, *p* < 0.05, one-way ANOVA; pairwise comparisons, WT Familiar versus WT Novel: *p* < 0.028, NTN1 cKO Familiar versus NTN1 cKO Novel: *p* < 0.96). All pairwise comparisons were performed with Tukey’s multiple comparison test. All data are shown as mean ± SEM.

### Selective deletion of netrin-1 from forebrain glutamatergic neurons reduces self grooming but does not elicit abnormal anxiety-like behaviour

Though widely used as a measure of spatial memory, the MWM has been reported to induce anxiogenic confounding behaviours due to its reliance on retrieval of emotionally aversive memories associated with the task (Harrison et al. 2009). Further, stress hormones can be elevated in rodents when assessed in the MWM, which may disturb or influence memory (Vogel-Ciernia and Wood 2014). To determine if mice lacking netrin-1 in excitatory neurons exhibited phenotypes that could impact performance in the MWM, we tested NTN1 cKO and age-matched littermate controls in an open field test to measure possible anxiety and motor abnormalities (Seibenhener and Wooten 2015). Analysis of the open field test relies on a rodent’s innate exploratory behaviour coupled with aversion to open spaces, and can be used to assess for hyper- or hypo-locomotor activity (Crawley 1985). A greater preference to travel along the boundary of the box compared to the center region is interpreted as increased anxiety. Mice were first habituated to the open field for 15 min before a subsequent 75 min experimental trial with movement trajectories recorded via an overhead camera (Figure 3A). Locomotor activity was expressed as distance travelled in successive 3 min bins over the course of the experimental trial (Figure 3B). No significant differences in locomotor activity were detected between NTN1 cKO and controls across the duration of the test. Additionally, no differences were observed between genotypes in the distance travelled along the border and center of the open field box (Figure 3C). Moreover, NTN1 cKO and controls did not differ in the total distance travelled (Figure 3D) or total activity counts (Figure 3E). These findings indicate that deletion of netrin-1 from glutamatergic neurons in the forebrain does not elicit gross-motor impairments or alteration of anxiogenic behaviours. This absence of changes in anxiety is also consistent with the lack of a difference between NTN1 cKO and control mice during the first three days of visible platform training in the MWM.

**Figure 3.**
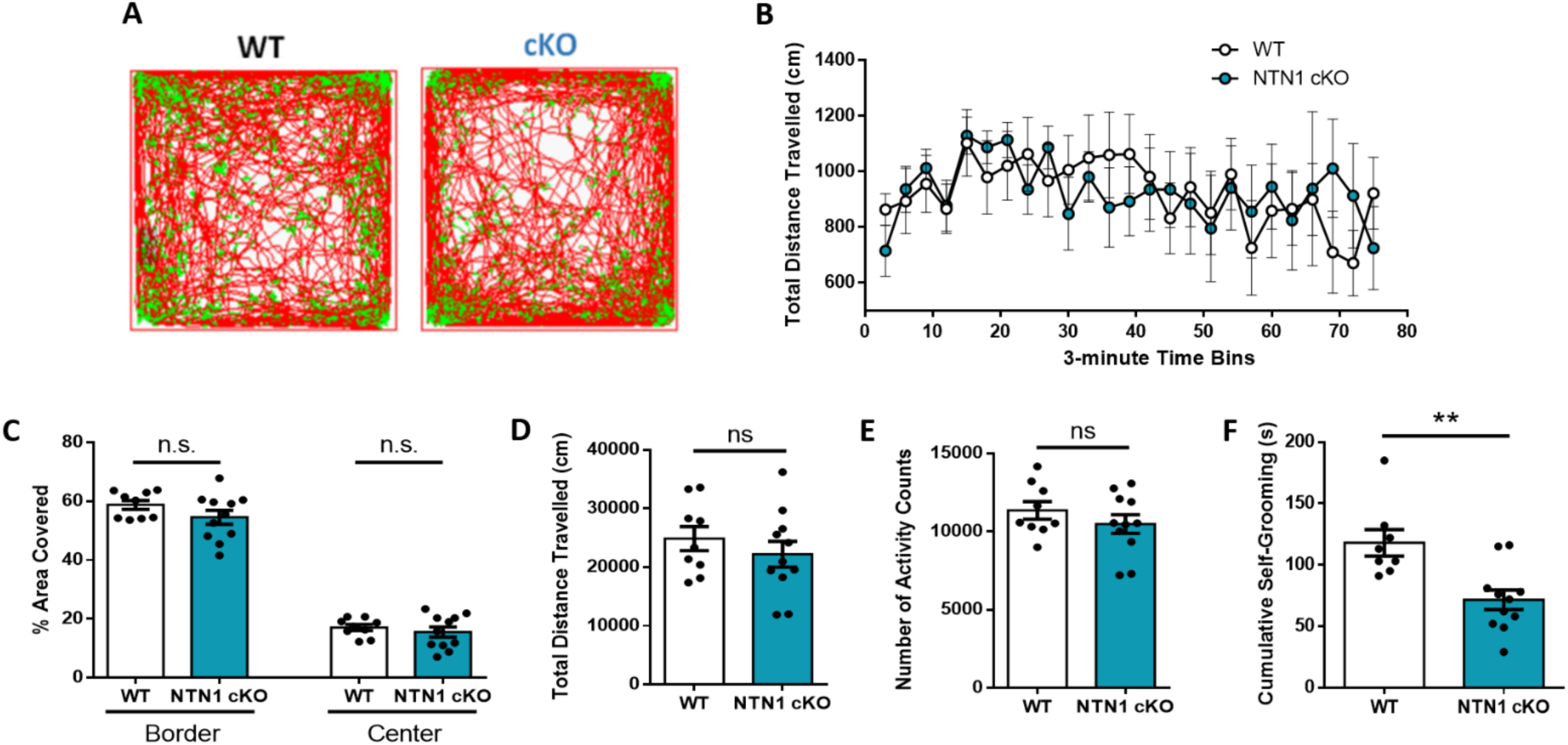
NTN1 cKO mice do not exhibit abnormal anxiety-like behaviours but show reduced self-grooming. (**A**) The trajectories travelled by the mice in the open field test were recorded by an overhead video-based tracking system. (**B**) Locomotor activity expressed as distance travelled in successive 3-min bins in the open field test (75-mins total). No differences were observed between wild-type and NTN1 cKO mice for all time bins (WT: *n* = 9, NTN1 cKO: *n* = 11, *p* > 0.05 for all time bins). (**C**) No significant differences were observed between genotypes for distance covered either within the border close to the walls of the chamber (WT: 58.75 ± 1.51, NTN1 cKO: 54.55 ± 2.38, *p* = 0.17) or within the center of the field (WT: 17.06 ± 1.05, NTN1 cKO: 15.49 ± 1.73, p = 0.47). Genotypes did not differ in the total distance travelled (**D**; WT: 24868 ± 2053, NTN1 cKO: 22217 ± 2190, *p* = 0.40) and the total counts of activity (**E**; WT: 11357 ± 558, NTN1 cKO: 10482 ± 594, *p* = 0.30) through the duration of the test. (**F**) NTN1 cKO mice displayed significant reduction in spontaneous self-grooming (WT: *n* = 8, 118.0 ± 10.72, NTN1 cKO: *n* = 11, 71.64 ± 7.97, p = 0.004). All comparisons were performed with two-tailed independent t-tests unless otherwise specified. Data are shown as mean ± SEM.

Grooming behaviour in rodents is an indirect measure of several behavioural phenomena such as motor sequencing and patterning, motivation, and anxiety (Kalueff et al. 2016) that are dependent on multiple brain regions including the limbic system and forebrain cortical regions. We therefore investigated whether the deletion of netrin-1 in NTN1 cKO mice might result in abnormalities related to repetitive grooming behaviours. Interestingly, NTN1 cKO mice displayed a significant reduction in the amount of time spent grooming relative to controls (Figure 3F).

### Selective deletion of netrin-1 from forebrain glutamatergic neurons does not impair novelty seeking behaviour

The NOPR test is commonly used as an alternative measure of hippocampal-dependent spatial memory that is less stressful than the MWM (Bannerman et al. 2014). However, due to its reliance on a rodent’s innate preference for novelty, a potential confounding aspect is that NTN1 cKO mice may exhibit deficits in novelty seeking behaviour, which may contribute to their poor performance in the NOPR task. To assess whether deletion of netrin-1 from forebrain glutamatergic neurons impairs novelty-seeking, we assessed NTN1 cKO and control age-matched wild-type littermates using a T-maze spontaneous alternation test (Figure 4A). This task is based on the innate motivation of rodents to explore novel environments. Mice were placed in the “starting arm” and allowed to choose a “goal arm” (ie. left or right arm). Following a 30 second delay, animals were placed back in the “starting arm” and again allowed to choose a “goal arm”. No significant difference was detected between genotypes in the spontaneous alternation task (Figure 4B), indicating that netrin-1 deletion in the NTN1 cKO mice does not disrupt preference for novelty.

**Figure 4.**
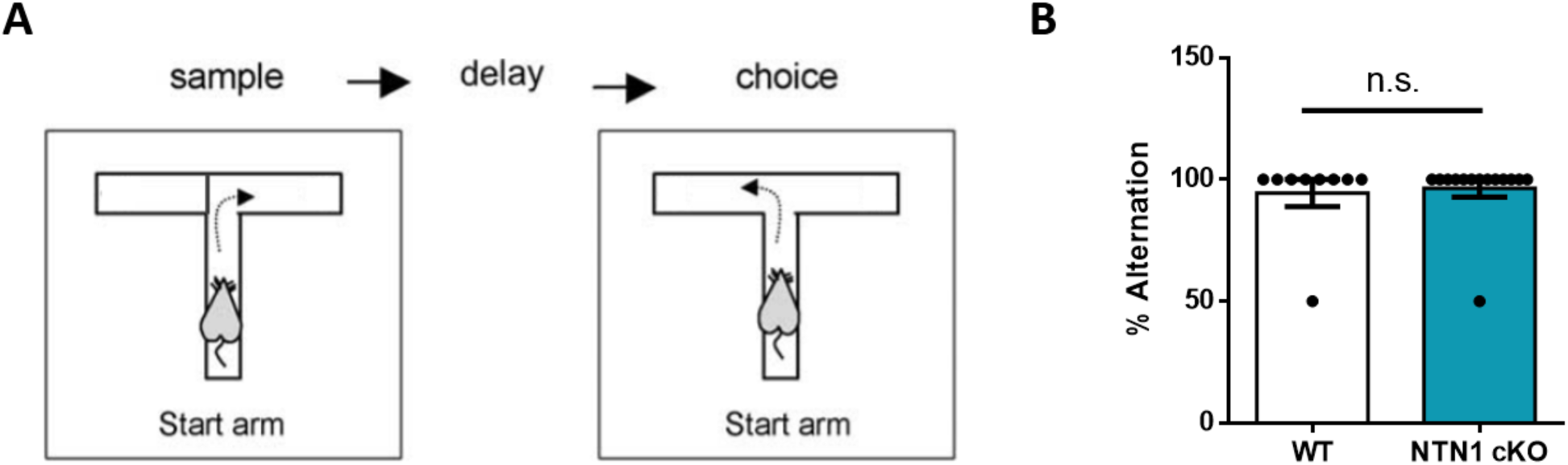
Mice with conditional forebrain netrin-1 deletion do not exhibit deficits in novelty preference. (A) Schematic representation of the spontaneous alternation T-maze test. (B) No differences were observed between age-matched wild-type and NTN1 cKO mice in the spontaneous alternation T-maze (WT: *n* = 9, 94.44 ± 5.56, NTN1 cKO: *n* = 14, 96.43 ± 3.57, *p* > 0.99, two-tailed Mann-Whitney test). Data shown as mean ± SEM).

## Discussion

The present study examined the behavioral consequences of conditional deletion of netrin-1 from forebrain glutamatergic neurons in adult mice. Previous studies indicate that netrin-1 and its receptor DCC are enriched at mature synapses, suggesting a role in synaptic transmission (Horn et al. 2013). Recent findings indicate that conditional deletion of netrin-1 from principal excitatory hippocampal neurons results in severe impairment in long-term potentiation (LTP) expression, and that netrin-1 secreted by neurons potently regulates synaptic transmission and plasticity in the adult hippocampus (Glasgow et al. 2018), a brain region critical for the consolidation of spatial memory (Nakazawa et al. 2004). Deficits in synaptic plasticity at Schaffer collateral synapses are associated with impairments in spatial memory retrieval and disruption of spatial representations (Moser et al. 1998; Brun et al. 2001). We hypothesized that deletion of netrin-1 expression in principal excitatory neurons would selectively impair spatial memory performance. To assess for a role in spatial memory, we tested the performance of mice conditionally-lacking netrin-1 expression in principal excitatory forebrain and hippocampal neurons (NTN1 cKO) in two hippocampal-dependent spatial memory tests: MWM and NOPR. Our findings demonstrated impaired spatial memory function in NTN1 cKO mice compared to age-matched control wild-type littermates. The observed deficits were specific to spatial memory, and no abnormalities in motor performance were detected in NTN1 cKO mice. Taken together, the present study reveals a critical role for netrin-1 expression by neurons in the regulation of the synaptic mechanisms that contribute to spatial memory.

### Conditional deletion of netrin-1 does not influence novelty-seeking

The CA3 region of the hippocampus has been proposed to serve as a comparator of novel stimuli, and hippocampal processing contributes to novelty detection in rodents (Vinogradova 2001; Kumaran and Maguire 2009). Disruption of the CA3 Schaffer collaterals synaptic inputs to CA1 pyramidal neurons impairs both spatial memory and novelty-seeking behaviour (Vago and Kesner 2008). Since the NOPR task relies on a rodent’s intrinsic preference for novelty, alterations in the mechanisms underlying novelty-seeking might contribute to impaired behavioural performance in NTN1 cKO mice. However, we observed no differences in spontaneous alternation on a T-maze task as an assessment of novelty-seeking. This indicates that loss of netrin-1 in forebrain glutamatergic neurons does not disrupt novelty detection, and supports the conclusion that the behavioral deficits observed are due to spatial memory dysfunction.

DCC haploinsufficient mice display blunted motor responses to amphetamine (Flores et al. 2005; Kim et al. 2013). These findings suggest that netrin-1 and DCC may influence the neural control of motor behaviour in rodents. While we did not detect a significant perturbation of locomotor activity in NTN1 cKO mice in the open field test or in the MWM, we did identify a decrease in spontaneous self-grooming by NTN1 cKO mice. Self-grooming is a highly evolutionary conserved behaviour that involves complex patterning of motor movements and is dependent on multiple brain regions, including the neocortex, striatum, and hypothalamus (Kalueff et al. 2016). Interestingly, disruption of glutamatergic synaptic activity in the neocortex has been reported to affect self-grooming (Aida et al. 2015; Kalueff et al. 2016). Although the specific neural mechanisms that underlie altered self-grooming remains unclear, enhanced cellular and synaptic excitability can increase repetitive behaviours. For example, mice lacking the astrocyte-specific glutamate transporter, GLT1, display aberrant excitatory transmission at corticostriatal synapses, along with increased self-grooming. In contrast, administration of an NMDAR antagonist, memantine, ameliorated the pathological self-grooming (Aida et al. 2015). We have previously demonstrated that netrin-1 alters NMDAR-dependent LTP, suggesting that deletion of netrin-1 may modify NMDAR function (Glasgow et al. 2018). As such, the observed reduced grooming may be a consequence of synaptic modulation mediated by NMDAR activation and changes in synaptic excitability due to a loss of netrin-1 expression.

### Netrin-1 and spatial memory consolidation

Netrin-1 signaling through DCC activates kinases involved in the regulation of LTP (Lai Wing Sun et al. 2011; Park et al. 2016; Glasgow et al. 2018; Incontro et al. 2018) and conditional deletion of netrin-1 or DCC from principal excitatory neurons severely attenuates an NMDAR-dependent form of LTP (Horn et al. 2013; Glasgow et al. 2018). In the developing nervous system, DCC directs cell motility through the activation of phospholipase C gamma, as well as regulating intracellular calcium, focal adhesion kinase, and local protein synthesis (Lai Wing Sun et al. 2011; Kang et al. 2018). NMDARs activate signaling pathways critical for LTP, including triggering the exocytosis of netrin-1 (Lynch et al. 1983; Glasgow et al. 2018). This local release of netrin-1 results in DCC-dependent and CaMKII-mediated recruitment of synaptic GluA1-containing AMPARs to facilitate synaptic transmission. Together, these findings indicate that netrin-1 signaling via DCC is critical to activate the mechanisms that underlie the long-term changes in synaptic strength associated with LTP.

Changes in synaptic strength are necessary for the formation of place cells, a subset of CA1 pyramidal neurons whose activity is linked with contextualized location-specific firing. Their activity is markedly elevated when an animal’s head is in specific regions of the environment (“place fields”) and virtually silent outside of these regions (O’Keefe and Dostrovsky 1971). The activity of a single place cell is correlated with cellular activity in adjacent place fields, which generates a spatial map of the environment (O’Neill et al. 2006). Importantly, replay of place cell activity between two synaptically-connected CA1 pyramidal neurons can induce NMDAR-dependent LTP (Isaac et al. 2009). In the intact animal, consolidation of spatial memory may be due to temporally-compressed replay of place cell sequences, resulting in high-frequency stimulation of synaptic inputs onto CA1 pyramidal neurons that triggers LTP-like synaptic consolidation (Nakazawa et al. 2004; Sadowski et al. 2016). Replay of place cell sequences is dependent on activation of NMDARs (Silva et al. 2015), which in turn can evoke netrin-1 release and activation of downstream signaling mechanisms involved in synaptic consolidation. Consequently, a lack of netrin-1 may disrupt the consolidation of spatial information; however, it remains unclear how netrin-1 might contribute to place cell formation and stabilization, as well as the network synchronization required for memory function.

Synchronous network activity plays a critical role in coordinating neuronal activity (Buzsaki 2002). Electrical field activity recorded in the hippocampus is dominated by theta-frequency (5-10 Hz) oscillations, large amplitude sinusoidal-like waveforms that are most prominent during periods of active exploration and rapid-eye movement (REM) sleep. Consistent with a critical role in synaptic plasticity, REM sleep theta activity and place cell replay are necessary to encode previously-acquired memories (Louie and Wilson 2001; Boyce et al. 2016). Oscillatory activity facilitates the coordination of synaptic inputs onto CA1 pyramidal neurons to increase postsynaptic depolarization (Montgomery et al. 2008). We predict this will promote netrin-1 exocytosis and thereby modify the synaptic connections that underlie the spatial memory; however, further studies are required to determine how netrin-1 modulates network-level activity and plasticity to influence the formation of spatial memories.

## Materials and Methods

### Animals

All procedures were performed in accordance with the Canadian Council on Animal Care guidelines for the use of animals in research and approved by the Montreal Neurological Institute Animal Care Committee. T29-1 CaMKIIα-Cre mice (Tsien et al. 1996) were obtained from The Jackson Laboratory (Bar Harbor, ME, USA), maintained on a C57BL/6 genetic background and crossed with mice homozygous for the floxed netrin-1 allele, netrin-1^*f/f*^ (NTN1^*f/f*^), also maintained on a C57BL/6 genetic background (Glasgow et al. 2018). *Cre* recombinase is first expressed ∼2.5 weeks postnatally, with expression throughout the forebrain but limited to glutamatergic neurons, including all hippocampal subfields by 1 month of age. Importantly, *cre* expression occurs well after the establishment of major axon tracts (Tsien et al. 1996; Horn et al. 2013). Previous work has demonstrated that netrin-1 protein levels in CaMKIIα-Cre-NTN1^*f/f*^ (NTN1 cKO) mice are significantly reduced by 3 months of age, therefore all experiments were performed with mice between 3 to 9 months old (Glasgow et al. 2018). We failed to observe significant effects of sex or age on behavioural measures, and therefore all data were pooled. Age-matched wild-type littermates were *cre-*negative NTN1^*f/f*^.

### Morris Water Maze

Spatial memory was evaluated in the Morris Water Maze (MWM), as described previously (Tong et al. 2012). NTN1 cKO and wild-type age-matched *cre-*negative littermates were trained on a modified 9-day protocol to assess for hippocampal-dependent spatial memory (Clark and Martin 2005). Briefly, mice were first trained for three days in a “visible” familiarization phase, during which a marked object was placed on the platform in a circular pool (140 cm diameter) filled with opaque, cold water (18±1°C), with visual cues located on the walls of the room equidistant above the water level. This was followed by five successive days of “hidden” platform testing where mice had to escape onto the platform relocated to a different quadrant within the pool and submerged ∼1 cm under the water surface, with spatial visual cues repositioned in the room. On day 1, animals that failed to locate the platform during the trial were guided to the platform and allowed to observe the visual cues for 10 secs. Mice that demonstrated consistent visual or motor abnormalities during the familiarization phase were removed from the analysis. Mice were randomly placed in a different area of the pool between training trials. Each trial lasted a maximum of 1 min. Twenty-four hours after the last training session, spatial memory was assessed using a probe trial in which the platform was removed. Escape latency during training, and automated unbiased analysis of movements during the probe trial, were measured using 2020 Plus tracking system and Water 2020 software (Ganz FC62D camera, HVS image).

### Novel Object Placement

Mice were trained and tested using the novel object placement recognition (NOPR) test as a non-invasive measure of hippocampal-dependent spatial memory, which lasted a total of 3 days (Boyce et al. 2016). On Day 1, mice were habituated to the square testing chamber (50 cm × 36 cm, 26 cm-high wall) for 5 mins without any added objects. On Day 2, during the “Sample Phase”, mice were exposed to two identical objects for 5 mins in two separate training sessions that took place 4 hours apart. Twenty-four hours following the last training session, one of the objects was moved to a novel location in the square chamber and the mice were provided 5 mins to explore both objects. An overhead camera recorded the mice in the square chamber throughout training and testing, and exploration time was measured by an experimenter blind to the genotypes. Objects and the test chamber were cleaned with 70% ethanol between trials to remove any olfactory cues.

Exploration times were calculated as the total time in the probe trial the animal investigated both objects (within one body length from an object with head pointed toward the object). Investigative ratios were calculated as the time spent exploring the novel placed object divided by the total time spent exploring both objects during the probe trial. Discrimination ratios were calculated as the difference between the time exploring the novel placed object and the time exploring the familiar placed object divided by the total time spent exploring both objects during the probe trial. Percentages of interactions with the novel placed object were reported as the number of contacts with the novel placed object divided by the total counts for both objects.

### Open Field Test

Mice were placed individually into clean open white square chambers (50 cm × 50 cm, 34 cm-high wall) for a total of 90 mins. Animal activity in the box was monitored using an infrared photobeam tracking system (VideoTrack, ViewPoint Life Sciences). Mice were habituated inside the open field for the first 15 mins, followed by the experimental trial, which lasted 75 mins (Seibenhener and Wooten 2015). Total distance travelled, number of activity counts (ie. initiation of movements), and time travelled were measured. The number of beam breaks were recorded every 3 mins. Image analysis of the distance spent in the border and center regions of the boxes was performed using *Fiji* software (Schindelin et al. 2012).

### Spontaneous Alternation T-Maze

The spontaneous alternation T-maze relies on a rodent’s attraction to explore novel environments and was used to assess for possible impairments in novelty-seeking behaviour (Deacon and Rawlins 2006). Mice were initially placed in the “start arm” of the T-maze (arms 30 cm × 10 cm, 20 cm high walls) and could subsequently choose a “goal arm” (Sample Phase). The animal was then confined within the chosen arm for 30 secs. A choice was defined as all four paws inside the arm. Following this, both the mouse and door were removed simultaneously, and the animal placed in the “start arm” facing away from the goal arms (Choice Phase). The animal was then allowed to again choose between the two open “goal arms” with the choice recorded. Based on the novelty of the previously unchosen arm, a correct choice occurs when the mouse alternates between arms, and an incorrect choice occurs when they animal does not alternate between arms. Two test trials, 24 hrs apart, were performed for each animal. For quantification, a score of either 100% or 0% was given to each mouse, and the average score for both trials per animal was calculated.

### Spontaneous Self-Grooming

Spontaneous self-grooming is commonly used as a measure of repetitive behaviour and motor coordination. Each mouse was individually placed in a novel mouse cage with a thin (1 cm) layer of bedding to reduce neophobia and prevent digging as a possible competing behaviour (Silverman et al. 2010). Following a 5 min habituation period in the test cage, each mouse was scored for the total time spent grooming all body regions for 10 mins (McFarlane et al. 2008). The observer sat approximately 2 m from the test cage and was blind to genotype.

### Statistical Analyses

Statistical analyses on parametric data were assessed using two-way repeated measures analysis of variance (ANOVA) followed by Bonferroni’s post-hoc test, one-way ANOVA followed by Tukey’s pairwise comparison test, and independent *t*-tests where appropriate. Analyses on non-parametric data were assessed using two-tailed Mann-Whitney test. Normality, homoscedasticity, and outlier tests were performed on all datasets. Statistical analyses were performed with GraphPad Prism. All data shown are presented as mean ± SEM (standard error of the mean), with statistical significance accepted as *p* < 0.05 using two-tailed tests.

## Author contributions

Conceptualization: EW, SDG, TEK; Methodology: EW, SDG, LJT; Investigation: EW, VR; Formal analysis: EW, SDG, LJT; Writing: EW, SDG, TEK; Funding: AS, ESR, EH, TEK; Resources, SDG, DC; Supervision: AS, ESR, EH, TEK

## Acknowledgments

The authors thank the members of the Kennedy lab for comments on drafts of the manuscript. We also thank Nathalie Marcal, Yi Jiang, Hanna Davies, Maumoud Moussa, and Hanan Elimam for technical assistance. EW was supported by a Healthy Brains for Healthy Lives (HBHL) scholarship. SDG was supported by postdoctoral fellowships from Fonds de la Recherche Québec – Santé (FRQS) and Canadian Institute for Health Research (CIHR). LJT was supported by scholarships from CIHR, Alzheimer Society of Canada, and Canadian Vascular Network. DC was supported by a scholarship from HBHL. The project was supported by grants from CIHR (AS, ESR, EH, TEK) and the Alzheimer Society of Canada (TEK). ESR holds a FRQS Research Chair.

